# Comparison of three air samplers for the collection of four nebulized respiratory viruses

**DOI:** 10.1101/2020.11.10.376053

**Authors:** Jasmin S Kutter, Dennis de Meulder, Theo M Bestebroer, Ard Mulders, Ron AM Fouchier, Sander Herfst

## Abstract

Viral respiratory tract infections are a leading cause of morbidity and mortality worldwide. Unfortunately, the transmission routes and shedding kinetics of respiratory viruses remain poorly understood. Air sampling techniques to quantify infectious viruses in the air are indispensable to improve intervention strategies to control and prevent spreading of respiratory viruses. Here, the collection of infectious virus with the six-stage Andersen cascade impactor was optimized with semi-solid gelatin as collection surface. Subsequently, the collection efficiency of the cascade impactor, the SKC BioSampler, and an in-house developed electrostatic precipitator was compared. In an in-vitro setup, influenza A virus, human metapneumovirus, parainfluenza virus type 3 and respiratory syncytial virus were nebulized and the amount of collected infectious virus and viral RNA was quantified with each air sampler. Whereas only low amounts of virus were collected using the electrostatic precipitator, high amounts were collected with the BioSampler and cascade impactor. The BioSampler allowed straight-forward sampling in liquid medium, whereas the more laborious cascade impactor allowed size fractionation of virus-containing particles. Depending on the research question, either the BioSampler or the cascade impactor can be applied in laboratory and field settings, such as hospitals to gain more insight into the transmission routes of respiratory viruses.

**Practical Implications:** Respiratory viruses pose a continuous health threat, especially to vulnerable groups such as young children, immunocompromised individuals and the elderly. It is important to understand via which routes these viruses can transmit to and between individuals that are at risk. If we can determine the amount of a certain respiratory virus in the air, then this will help to predict the importance of transmission through the air for this virus. Most currently available air sampling devices have not been designed to collect infectious viruses from the air. Therefore, we here optimized and compared the performance of three air samplers for four different respiratory viruses.

## Introduction

Viral respiratory tract infections occur frequently and are a leading cause of morbidity and mortality worldwide^1^. Despite their impact on public health, for most respiratory viruses little is known about the relative contribution of various routes of transmission. Most of the current knowledge of respiratory virus transmission has been derived from experimental studies (e.g. human challenge studies, animal transmission experiments and virus stability studies) and observational epidemiological studies during outbreaks^2^.

Respiratory viruses can spread via different routes: direct contact (e.g. via hand shaking), indirect contact (via contaminated surfaces) or through the air via droplets and/or aerosols. We define droplets as particles that can only travel short distances through the air before they settle onto the mucosa of individuals or nearby surfaces, and aerosols as particles that are small enough to remain suspended in the air for prolonged periods of time and cover large distances. To understand the relative importance of various transmission routes, knowledge on the viral load and infectivity of viruses in the air, as well as the size of virus-containing droplets and aerosols is warranted. To meet this need, the collection of viruses from the air with air samplers has gained increasing attention. Such air samplers can be applied in environments such as hospital settings, animal experiments or livestock farms^3–6^, however, collecting infectious virus from air remains challenging^7,8^. A limitation is that air samplers have a high cut-off size, which prevents efficient collection of small aerosols (<1 μm). In addition, the high collection forces and sampling velocities applied inside the air samplers can damage virus particles, resulting in the collection of mainly non-infectious virus^4,9^.

In only a few studies, infectious respiratory viruses were collected from air as demonstrated by virus isolation in cell cultures^10–13^, whereas in most studies the presence of virus in air was solely determined by the detection of viral RNA by (quantitative) RT-PCR^14–16^.

Here, existing and newly developed air samplers with different collection methods were improved and compared: the six-stage Andersen cascade impactor, the SKC BioSampler and an in-house developed electrostatic precipitator. The BioSampler and cascade impactor are commercially available air samplers that employ inertial forces to remove aerosols and droplets from the air flow. The cascade impactor was originally developed to collect airborne bacteria and fungi onto petri dishes filled with bacteriological agar. However, agar is less suitable as collection medium for viruses because of the high impaction forces and desiccation effects on aerosols and droplets^17–19^.

For the cascade impactor and BioSampler, the collection efficiency is low for aerosols in the submicron range^20–22^. To overcome the poor collection efficiency of small aerosols, we developed an electrostatic precipitator. Electrostatic precipitators are widely used to remove small particles such as dust from air, and they are being increasingly explored for air sampling of airborne microorganisms^23–26^. In electrostatic precipitators, the air around a conductor is ionized through the application of high voltage. Incoming droplets and aerosols get charged and attracted into a neutral or oppositely charged collection medium. These air samplers have a low flow rate and subject aerosols and droplets to less physical stress, thereby yielding higher recovery rates of infectious microorganisms^25,26^.

In this study the collection of infectious virus with the cascade impactor was first improved by optimizing the collection medium. Subsequently, the efficiency to collect infectious virus and viral RNA of the BioSampler, cascade impactor and electrostatic precipitator for nebulized pandemic H1N1 influenza A virus (pH1N1), human metapneumovirus (HMPV), human parainfluenza virus type 3 (PIV3) and respiratory syncytial virus (RSV) was compared in an in-vitro setup. Finally, the sensitivity of the BioSampler and cascade impactor for low virus concentrations was evaluated.

## Material and Methods

### Viruses

Human H1N1 influenza A virus A/Netherlands/602/2009 (pH1N1) was propagated in Madin-Darby canine kidney (MDCK) cells. Recombinant HMPV NL/1/00 expressing green fluorescent protein (GFP) and GFP-expressing PIV3 (ViraTree) were propagated in subclone 118 of Vero-WHO cells (Vero-118 cells)^27,28^. RSV A2 (ATCC) was propagated in human epithelial 2 (Hep-2) cells.

### Cells

MDCK cells (ATCC) were cultured in Eagle’s minimal essential medium (EMEM; Lonza) supplemented with 10% fetal bovine serum (FBS; Greiner or Atlanta Biologicals), 100 IU/ml penicillin-100 μg/ml streptomycin mixture (Lonza), 2 mM L-glutamine (Lonza), 1.5 mg/ml sodium bicarbonate (Lonza), 10 mM Hepes (Lonza) and 1x nonessential amino acids (Lonza). Vero-118 cells were cultured in Iscove’s Modified Dulbecco’s Medium (IMDM) supplemented with 10% FBS, 100 IU/ml penicillin-100 μg/ml streptomycin mixture (Lonza) and 2 mM L-glutamine (Lonza). Hep-2 cells were cultured in Dulbecco’s Modified Eagle Medium (DMEM; Lonza or Gibco) supplemented with 10% FBS; Greiner or Atlanta Biologicals), 100 IU/ml penicillin-100 μg/ml streptomycin mixture (Lonza), 2 mM L-glutamine (Lonza), 1.5 mg/ml sodium bicarbonate (Lonza), 10 mM Hepes (Lonza) and 0.25 mg/ml fungizone (Invitrogen). All cells were cultured at 37°C and 5% CO2.

### Virus titrations

For endpoint titration of viruses, cells were grown to confluency in 96 well plates overnight. Subsequently, cells were inoculated with 100 μl of 10-fold serial dilutions of collected air samples or controls. One hour after inoculation, cells were washed once and cultured in infection medium consisting of either serum free EMEM supplemented with 100 IU/ml penicillin-100 μg/ml streptomycin mixture (Lonza), 2 mM L-glutamine (Lonza), 1.5 mg/ml sodium bicarbonate (Lonza), 10 mM Hepes (Lonza) and 1x nonessential amino acids (Lonza) and 20 μg/ml *N*-tosyl-l-phenylalanine chloromethyl ketone (TPCK) treated trypsin (Sigma Aldrich) for MDCK cells, serum-free IMDM supplemented with 100 IU/ml penicillin-100 μg/ml streptomycin mixture (Lonza) and 2 mM L-glutamine (Lonza) and 3.75 μg/ml trypsin (BioWhittaker) for Vero-118 cells and serum reduced (2%) DMEM supplemented with 100 IU/ml penicillin-100 μg/ml streptomycin mixture (Lonza), 2 mM L-glutamine (Lonza), 1.5 mg/ml sodium bicarbonate (Lonza), 10 mM Hepes (Lonza) and 0.25 mg/ml fungizone (Invitrogen) for Hep-2 cells. For pH1N1 virus, supernatants of cell cultures were tested for agglutination activity using turkey erythrocytes after 3 days of incubation. For RSV, cell cultures were observed for CPE after 7 days of incubation. For GFP-expressing HMPV and PIV3, wells were screened for GFP positive cells using an inverted fluorescence microscope at 7 and 5 days of incubation, respectively. Infectious virus titers were calculated from four replicates as tissue culture infective dose (TCID_50_) by the Spearman-Karber method.

### Real-time quantitative RT-PCR

Viral RNA was extracted from 200 μl sample and eluted in a total volume of 50 μl using the MagNA Pure LC Total Nucleic Acid Isolation Kit, according to instructions of the manufacturer (Roche). Twenty μl of virus RNA was amplified in a final volume of 30 μl, containing 7,5 μl 4xTaqMan Fast Virus 1-Step Master Mix (Life Technologies) and 1 μl Primer/Probe mixture^29,30^. Amplification was performed using the following protocol: 5 min 50°C, 20 sec 95°C, 45 cycles of 3 sec 95°C and 31 sec 60°C.

### Air samplers

The six stage Andersen cascade impactor (Thermo Scientific) operates at 28.3 liters per minute (LPM) and consists of six stages, with 400 orifices each, and 6 petri dishes^31^ (**Fig 1A**). With increasing stage number, the size of the orifices decreases and hence impaction velocity increases, enabling the collection of size-fractionated droplets and aerosols over the different petri dishes. Based on solid impaction, bacteria and fungi were originally captured onto petri dishes filled with bacteriological agar. For virus collection, virus transport medium (VTM) and an in-house developed semi-solid gelatin layer were compared with the conventional agar. VTM consisted of Minimum Essential Medium (MEM) – Eagle with Hank’s BSS and 25 mM Hepes (Lonza), glycerol 99% (Sigma Aldrich), lactalbumin hydrosylate (Sigma Aldrich), 10 MU polymyxin B sulphate (Sigma Aldrich), 5 MU nystatin (Sigma Aldrich), 50 mg/ml gentamicin (Gibco) and 100 IU/ml penicillin 100 μg/ml streptomycin mixture (Lonza), while the semi-solid gelatin layer was prepared from commercial gelatin sheets (Dr. Oetker) dissolved in VTM (10 mg/ml). For all collection media, polystyrene 100 mm petri dishes (Greiner) were used. To maintain an optimal jet-to-plate distance with the polystyrene dishes, a total volume of 41 ml was used to fill the plates based on manufacturer instructions. To avoid high dilution factors of the samples, petri dishes were first filled with 32 ml of 2% agarose (Roche) as a bottom layer on which the actual collection medium, 9 ml VTM, or 9 ml semi-solid gelatin, was placed. For the agar impaction surface, 41 ml of 1.5% w/v bacteriological agar NO.1 (Thermo Scientific™) was used. To quantify collected infectious virus and total virus RNA, samples were processed in liquid form. Semi-solid gelatin was liquefied directly after sampling by adding 6 ml of prewarmed (37°C) VTM to each plate followed by incubation for 30 min at 37°C. Agar plates were carefully scraped with a cell scraper after adding 6 ml VTM. VTM samples were simply aspirated from the petri dishes. Samples were aliquoted and titrated or stored at −80°C for subsequent RNA isolation and qRT-PCR analysis.

**Figure 1.**
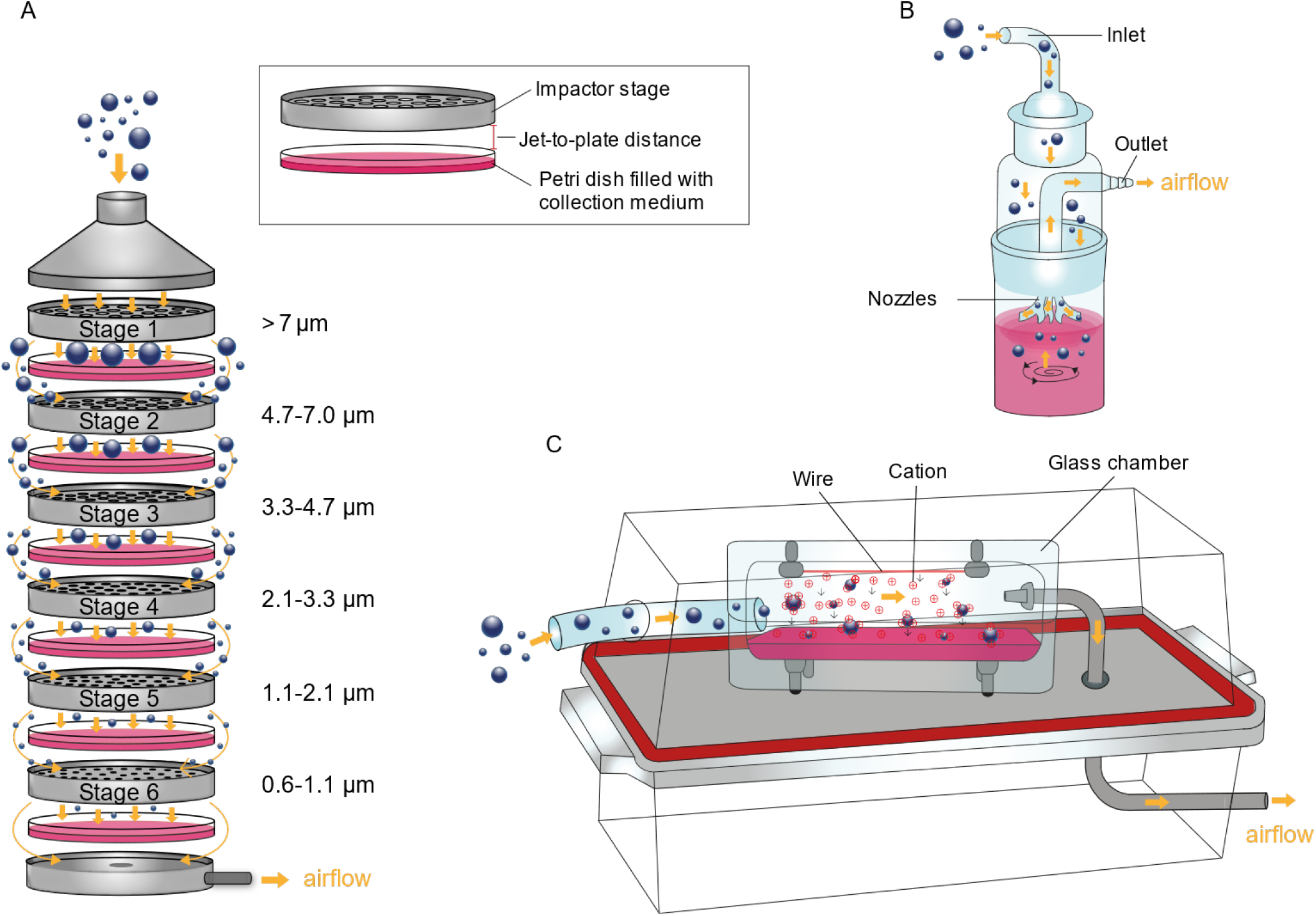
The different samplers that were compared in this study. A. Six-stage Andersen cascade impactor. Aerosols and droplets are collected from the air according to their size in 10cm dishes filled with semi-solid gelatin, agar or VTM. An accurate jet-to-plate distance is important to ensure a correct size-fractionation. B. SKC BioSampler (all-glass impinger). Air is drawn in and accelerated in the three nozzles. Particles are subsequently collected into swirling VTM by impingement. C. Electrostatic precipitator. Inside a Nalgene box, air is drawn into a glass chamber in which air is ionized. Cations bind to the particles and drag aerosols and droplets to the bottom reservoir which is filled with VTM. Orange arrows indicate air flow. Blue spheres indicate aerosols and droplets of different sizes.

The SKC BioSampler (SKC Inc) is an all-glass impinger that utilizes a liquid collection medium to capture droplets and aerosols (**Fig. 1B**). It consists of an inlet, a collection vessel and an outlet. The inlet contains three 0.63 mm tangential nozzles through which air is drawn at a flow rate of 12.5 LPM, thereby creating a swirling motion in the liquid collection medium. The swirling motion minimizes the chances of particle re-nebulization and maintains the infectiousness of collected particles. When aerosols and droplets exit the nozzles they get impinged into the liquid medium, while the remaining air exits the air sampler through the outlet. As collection medium, 15 ml virus transport medium (VTM) was used. Twenty μl antifoam B emulsion (Sigma Aldrich) was added to prevent the generation of bubbles and foam due to the swirling motion of the collection medium during air sampling.

The in-house developed electrostatic precipitator is made of a glass chamber consisting of an upper and bottom part (**Fig 1C**). A voltage of 13 kV is applied to a 80 mm long corona wire that is attached to the upper part of the glass chamber with a distance of 20 mm between the wire and the bottom part. Application of high voltage produces an ion discharge which ionizes the air in the chamber. Upon collision of incoming aerosols and droplets with ionized air molecules, aerosols and droplets become charged and attracted by the neutral bottom part of the glass chamber which is filled with 20 ml VTM. As a side effect, corona discharges also generate ozone, which is known to inactive viruses^32,33^. A positive charge was used in the electrostatic precipitator because it produces less ozone than a negative charge^34,35^. The electrostatic precipitator is operated at 4 LPM.

### Experimental air sampling setup

Air samplers were connected to a Nalgene BioTransport Carrier box (dimensions 36,8 x 18,4 x 17,0 cm (L x W x H)), in which 500 μl of a virus suspension was nebulized using the Aerogen Solo nebulizer (Medicare Uitgeest B.V.). The vacuum pump was switched on just before nebulization and air was drawn through the air samplers for a total of 5 minutes **(Fig 2).** Subsequently, air samples were retrieved from the samplers, agar and semi-solid gelatin samples were processed as described above and all samples were subjected to further analysis. All experiments were performed in a class 2 biosafety cabinet.

**Figure 2.**
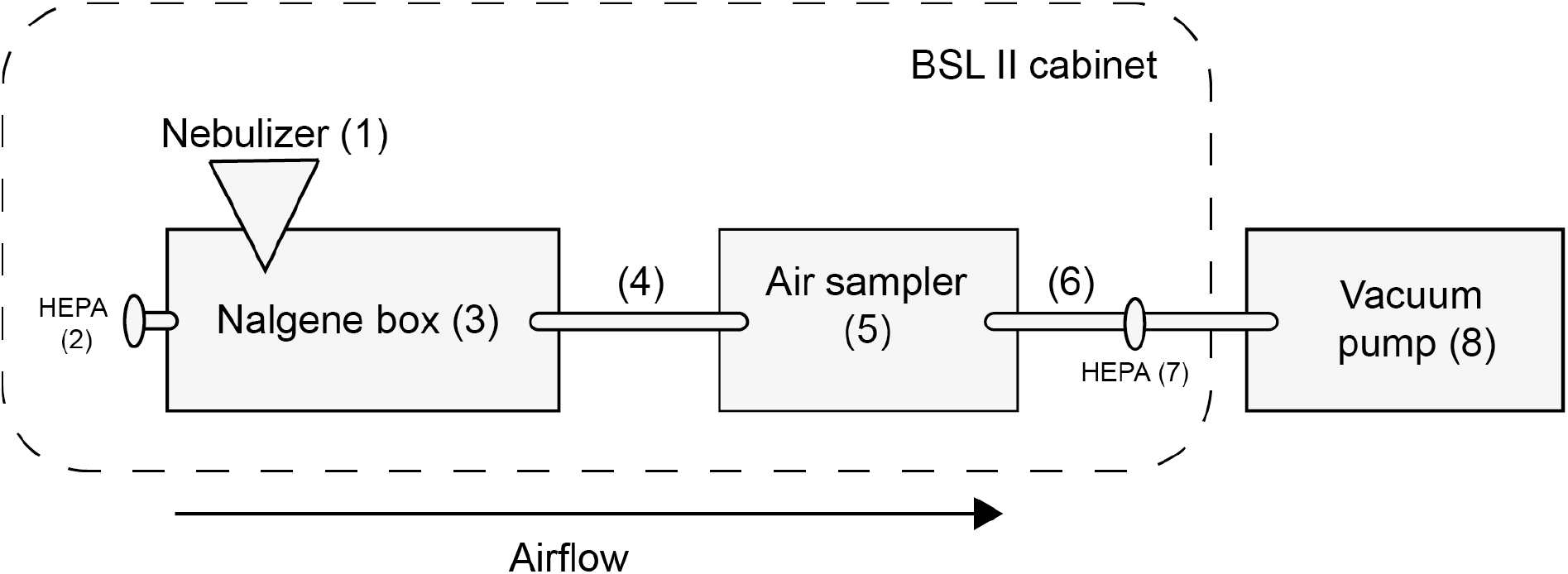
Schematic representation of the experimental air sampling setup. Virus suspensions were nebulized using a nebulizer (1) to generate aerosols and droplets containing virus particles in a Nalgene box (3), which was connected via a tube (4) to an air sampler (5). A second tube (6) was placed between the air sampler and a vacuum pump (8) that was placed outside the BSL II cabinet. High-efficiency particulate air (HEPA) filters (2,7) were installed on both sides of the air sampling set-up to guarantee that clean air entered the box and to prevent contamination of the environment. For each experiment, nebulized viruses were collected from the Nalgene box with air samplers for 5 min.

## Results

### Loss of virus infectivity due to mechanical nebulization

Since virus might lose infectivity during the mechanical nebulization of virus suspensions using the Aerogen Solo nebulizer, the loss of virus infectivity during this process was first quantified. For this purpose, 500 μl of pH1N1 virus, HMPV, PIV3 or RSV was either directly pipetted into 15 ml of VTM in a 50 ml tube (positive control), or nebulized into 15 ml VTM in a T75 cell culture flask. A direct comparison of the titers with and without nebulization demonstrated that pH1N1 virus infectivity was barely affected by this process, as the virus titer after nebulization was only 0.03 log_10_TCID_50_ lower than without nebulization. The virus titers of HMPV, PIV3 and RSV were reduced by 0.58, 0.55 and 0.54 log_10_TCID_50_, respectively **(Table 1)**. The loss of virus infectivity during nebulization may be due to incomplete nebulization of virus suspensions or due to viruses not being resistant to the mechanical forces applied during nebulization. Overall, the loss of virus infectivity due to nebulization was only marginal, hence this method was used in subsequent experiments.

**Table 1.** Loss of virus infectivity due nebulization with the Aerogen solo nebulizer.

| **Virus** | **Virus titer before**<br>**nebulization**<br>**(log<sub>10</sub>TCID<sub>50</sub> +/-SD)<sup>†</sup>** | **Virus titer after**<br>**nebulization**<br>**(log<sub>10</sub>TCID<sub>50</sub> +/-SD)<sup>†</sup>** |
| --- | --- | --- |
| **pH1N1** | 6.36 +/- 0.46 | 6.33 +/- 0.58 |
| **HMPV** | 6.20 +/- 0.7 | 5.62 +/- 0.73 |
| **PIV3** | 6.53 +/- 0.3 | 5.98 +/- 0.2 |
| **RSV** | 6.86 +/- 0.16 | 6.32 +/- 0.43 |

<sup>†</sup>Average titers of six replicates

### Optimization of the virus collection efficiency of the cascade impactor

Since agar was expected to be less suitable as collection medium for viruses, two other collection media were tested in addition to agar: an in-house developed semi-solid gelatin layer and VTM. High doses of pH1N1 virus or HMPV were nebulized into the Nalgene box and subsequently collected from the air using the cascade impactor containing petri dishes filled with either agar, semi-solid gelatin or VTM. For the dishes filled with agar, 6 ml VTM was added followed by careful scraping with a cell scraper. For the petri dishes filled with semi-solid gelatin, 6 ml prewarmed (37°C) VTM was added followed by incubation for 30 min at 37°C to liquefy the semi-solid gelatin. When only VTM was used as collection medium, the medium was simply collected from the petri dishes after sampling. After this post-sampling processing, samples from all six stages were subjected to virus titration and qRT-PCR to determine the amount of infectious virus and viral RNA collected in each stage. Subsequently, the total amount (i.e. the sum of collected virus of all six stages) of infectious virus and viral RNA was calculated and compared to that of the positive control, which was 15 ml of VTM containing the same amount of virus as was nebulized and collected by the air sampler. For pH1N1 virus, collection of infectious virus was equally efficient when agar or semi-solid gelatin was used, and only 0.6 and 0.8 log_10_TCID_50_ were lost, respectively, as compared to the positive control. Collection of infectious pH1N1 virus in VTM was much less efficient and resulted in a reduction of 2.4 log_10_TCID_50_ **(Fig 3A)**. The total collection efficiency of the cascade impactor for pH1N1 virus RNA as compared to the positive control was 16.4% and 6.4% for agar and semi-solid gelatin, but interestingly 39.4% for VTM, despite the substantial loss of virus infectivity **(Fig 3B).** Also, for HMPV, the amount of infectious virus collected with each medium varied. The collection efficiency was highest with semi-solid gelatin and VTM, where 1.4 and 1.6 log_10_TCID_50_ less infectious virus was recovered after air sampling as compared to the positive control **(Fig 3C)**. When agar was used as collection medium, considerably less infectious HMPV was collected with a reduction of 2.4 log_10_TCID_50_ as compared to the positive control **(Fig 3C)**. The total collection efficiency of the cascade impactor for HMPV RNA varied from 1.4%, 5.4% and 12.3% for semi-solid gelatin, agar and VTM respectively. Thus, also for HMPV the highest physical collection efficiency was obtained with VTM. **(Fig 3D)**.

**Figure 3.**
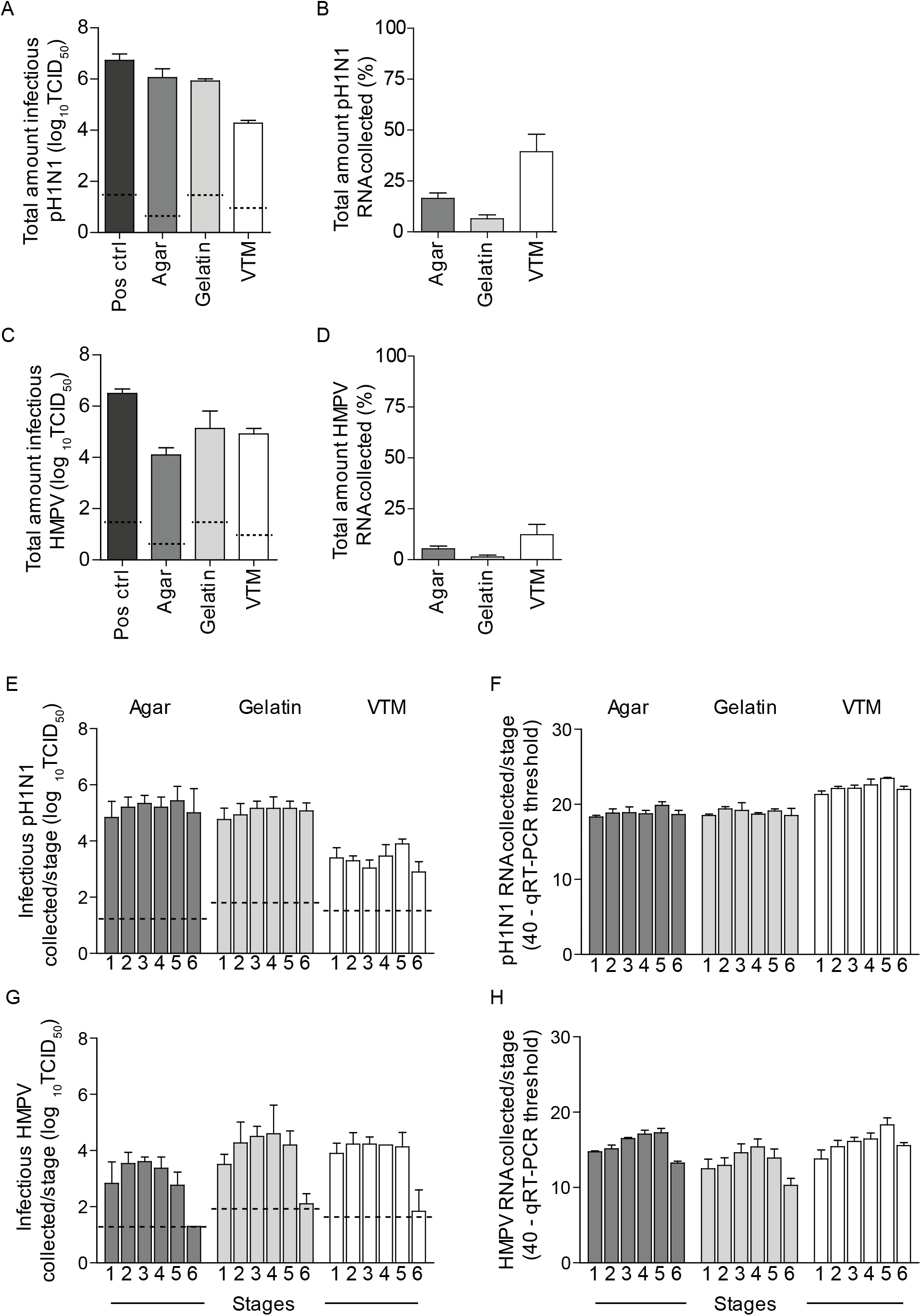
Evaluation of different collection media for the cascade impactor. pH1N1 virus and HMPV were collected on agar, semi-solid gelatin or VTM to compare the collection efficiency of the cascade impactor with each medium. For both viruses and the different collection media, the total amount of collected infectious virus (A and C) and viral RNA (B and D) as well as the distribution of the amount of infectious virus (E and G) and viral RNA (F and H) over the six stages is shown. Dotted lines indicate the detection limit of the virus titrations. Bars represent mean values of 3 experiments. Error bars indicate SD of 3 experiments.

To size fractionate virus-containing particles in the cascade impactor, the air velocity and impaction forces increase with increasing stage number. As a consequence, viruses may be subjected to more physical stress in stage 6 as compared to stage 1. Therefore, to investigate if the virus infectivity was differently conserved over all stages, the amounts of infectious virus and virus RNA collected in the individual stages, were also compared. When agar, semi-solid gelatin or VTM was used to collect pH1N1 virus, the amounts of infectious virus and virus RNA were evenly distributed over all stages, suggesting that the infectivity of pH1N1 virus was not more affected in the higher stage numbers as compared to the lower stage numbers. **(Fig 3E and 3F)**. The distribution of infectious HMPV and HMPV RNA over the six stages was slightly more variable than that of pH1N1 virus. Interestingly, substantially lower amounts of infectious HMPV and viral RNA were captured in stage 6 as compared to pH1N1 **(Fig 3G and 3H)**. Overall, the infectivity of both viruses was well conserved with semi-solid gelatin, which was therefore used in subsequent experiments.

### Comparison of the collection efficiency of three air samplers for four common respiratory viruses

After improving the collection of infectious viruses from the air with the cascade impactor, the collection efficiency of the BioSampler, the cascade impactor with semi-solid gelatin as collection medium and the electrostatic precipitator was assessed with four different respiratory viruses. The three air samplers employ different collection methods and use different flow rates and collection media, and can thus differ in their ability to efficiently collect viruses from air. In addition, the collection efficiency may not be the same for all respiratory viruses, as their stability in air and sensitivity to the mechanical forces applied in the air samplers may vary. High doses of pH1N1 virus, HMPV, PIV3 and RSV were nebulized and subsequently collected with each air sampler. The highest collection efficiency of infectious virus was obtained with the cascade impactor for pH1N1 virus and the BioSampler for HMPV and RSV, whereas similar amounts of PIV3 were collected with the cascade impactor and BioSampler **(Fig 4A)**. For all four viruses only low amounts of infectious virus were collected with the in-house developed electrostatic precipitator **(Fig 4A)**. The collection efficiency of the electrostatic precipitator for virus RNA was also very low demonstrating that the overall collection of viruses with this sampler was poor **(Fig 4B)**.

**Figure 4.**
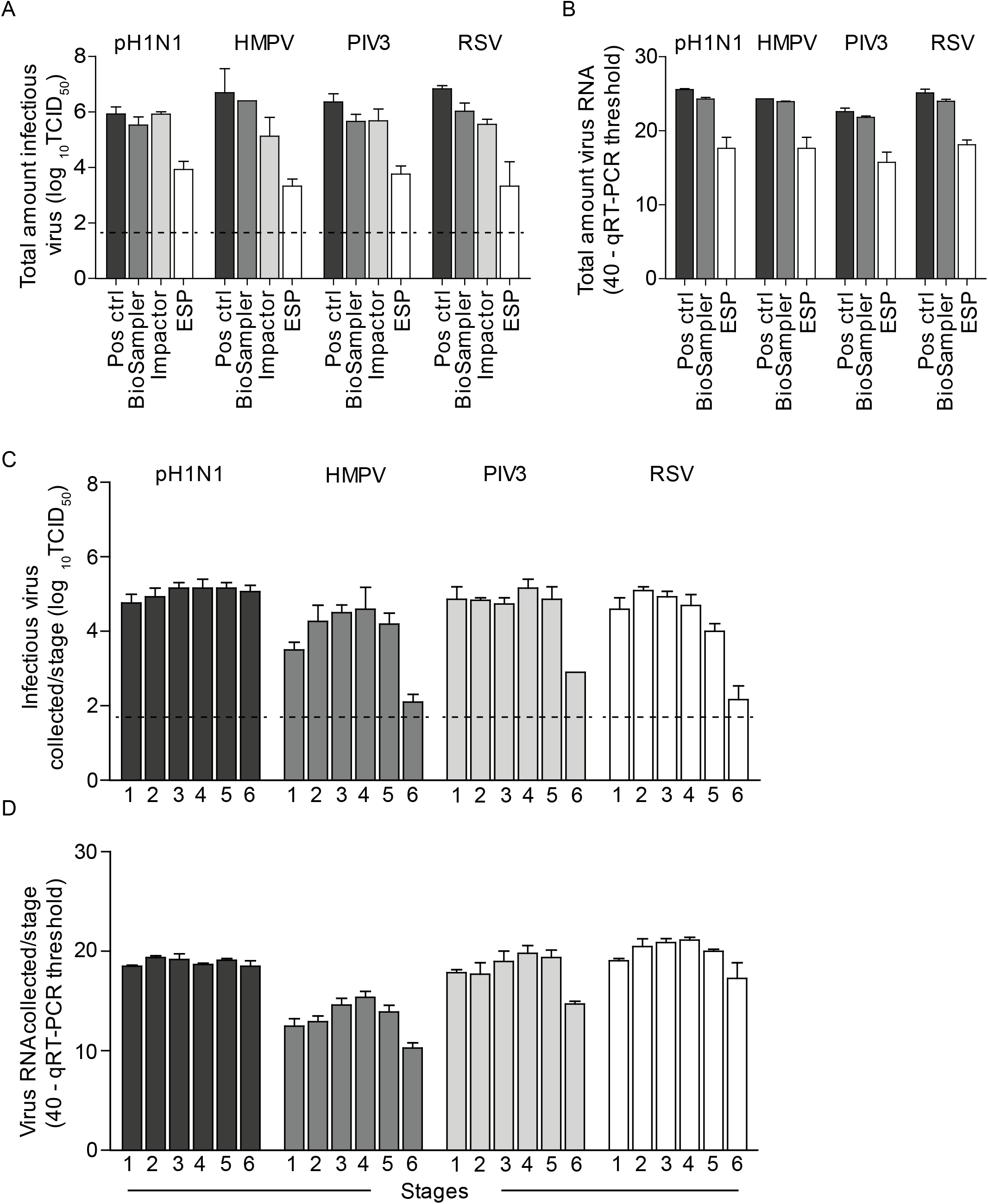
Performance of all air samplers with different respiratory viruses. To compare the performance of the three air samplers, pH1N1 virus, HMPV, PIV3 and RSV were each nebulized and collected with the BioSampler, cascade impactor (with semi-solid gelatin) and electrostatic precipitator. For all viruses, the amount of collected infectious virus (A and C) and viral RNA (B and D) is shown for each air sampler. Dotted lines indicate the detection limit of the virus titrations. Bars represent mean values of 3 experiments. Error bars indicate SD of 3 experiments.

When the distribution patterns of the four viruses over the six stages of the cascade impactor were investigated, the amounts of collected infectious pH1N1 virus and pH1N1 RNA was found to be similar in all stages. In contrast considerably lower amounts of infectious virus and virus RNA was collected in stage 6 for HMPV, PIV3 and RSV as compared to the other stages **(Fig 4C and 4D)**.

### Sensitivity of the BioSampler and the cascade impactor

An ideal air sampler is capable of collecting small amounts of infectious virus from a large air volume in a small sample volume. Therefore, the sensitivity for collecting infectious viruses from the air was assessed for the BioSampler and cascade impactor, the two samplers with the highest collection efficiency in this study. Approximately 10^5.7^ and 10^3.7^ TCIDδ¤ of pH1N1 virus and HMPV were nebulized and collected as described above. Despite the lower amounts of nebulized virus, both air samplers were still able to collect infectious virus as efficient as when high amounts virus were nebulized. Air sampling with the BioSampler resulted in a reduction of 0.2 and 0.3 log_10_TCID_50_ for pH1N1 virus, and 1.4 and 1.0 log_10_TCID_50_ for HMPV, as compared to the positive control, when 10^5.7^ and 10^3.7^ TCID_50_ of virus was nebulized, respectively **(Fig. 5A and 5B)**. The cascade impactor collected 0.6 and 0.8 log_10_TCID_50_ less pH1N1 virus, and 0.9 and 0.9 log_10_TCID_50_ less HMPV, as compared to the positive control, when 10^5.7^ and 10^3.7^ TCID_50_ were nebulized, respectively **(Fig. 5A and 5B)**.

**Figure 5.**
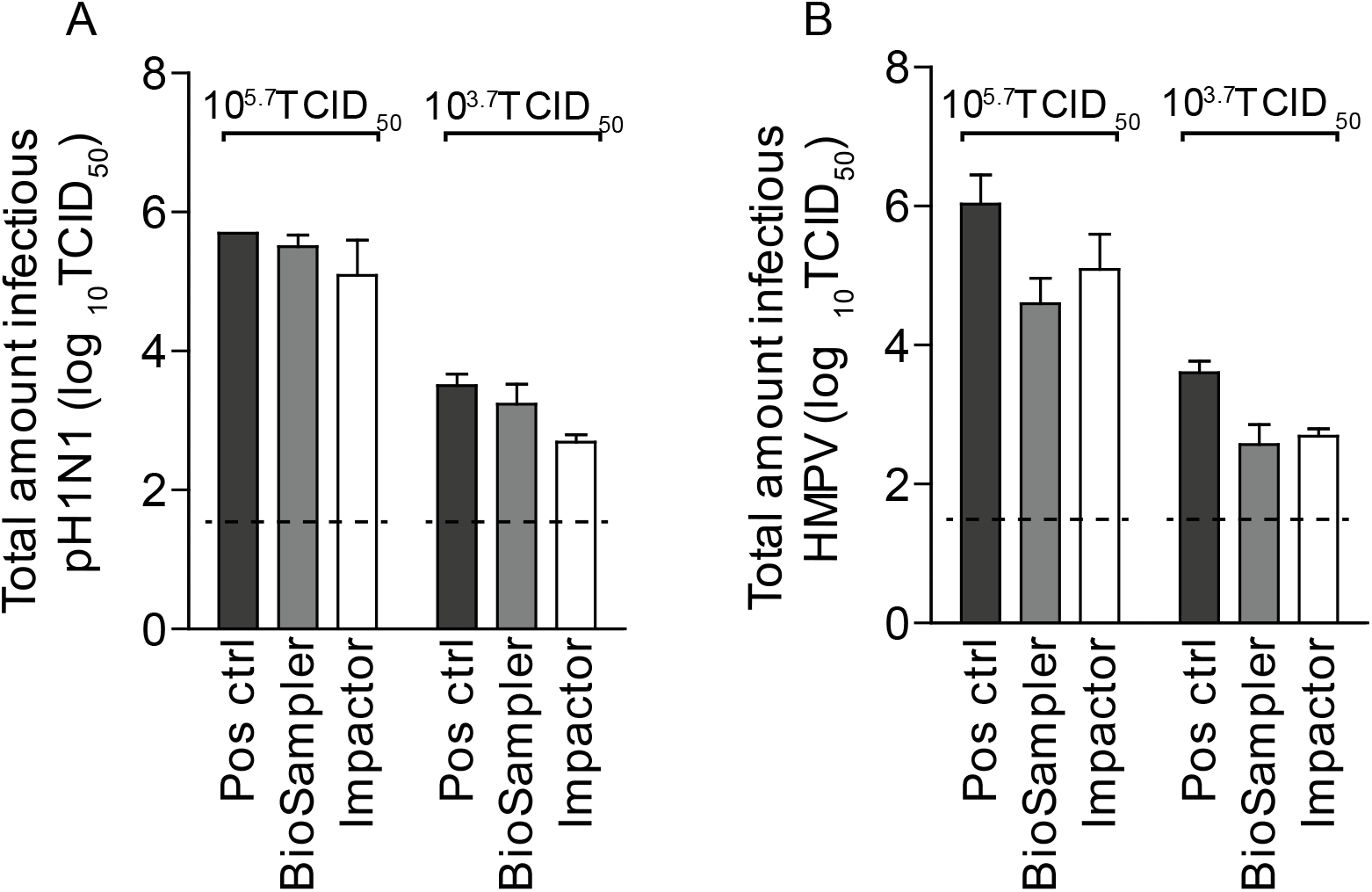
Collection efficiency of the BioSampler and cascade impactor for two lower doses of infectious virus. 10^5.7^ and 10^3.7^ TCID_50_ of pH1N1 virus (A) and HMPV (B) were nebulized and the total amount of infectious virus was determined by virus titration. Dotted lines indicate the detection limit of virus titrations. Bars represent mean values of 3 experiments. Error bars indicate SD of 3 experiments.

## Discussion

Air sampling is increasingly recognized as an important tool for the characterization and quantification of respiratory viruses in the air in different environments, such as hospital settings, epidemiological investigations and laboratory experiments. Information on the amount of infectious virus in the air, the ability of a virus to remain infectious in the air and the size distribution of droplets and aerosols that contain infectious viruses will help to identify the relative contribution of the possible transmission routes of the respiratory virus under investigation. For this purpose, here, the performance of three air samplers which employ different collection methods and use different collection media was compared in an in-vitro setup by evaluating their efficiency to collect four artificially nebulized respiratory viruses.

In cascade impactors, originally agar was used to collect bacteria from air. However, agar is generally considered less suitable as a collection surface for viruses, given the possibility of desiccation and increased particle bounce^36–38^. In addition, it was demonstrated for infectious bursal disease virus, that virus recovery from agar is reduced significantly when petri dishes are processed at later time points, and immediate processing is not always possible in field studies^39^.

As an alternative to agar, liquid medium is also frequently used in the cascade impactor, since the chances of virus desiccation are smaller and sample processing after collection is not needed^15,40–42^. However, the high flow velocities within the sampler push aside liquid medium where the air stream hits the surface, creating a dent, thereby increasing the jet-to-plate distance. This may result in a shift of size fractionation, with larger particles being collected in lower stages and smaller particles escaping from collection by the cascade impactor. Furthermore, liquid spill-over into other stages increases the chances of cross-contamination and VTM can be spilled easily, making the transport of petri dishes challenging, as also reported by others. The results of the present study show that the collection of infectious pH1N1 virus in agar was more efficient than in VTM, while the opposite was true for HMPV, indicating that different collection media may be required depending on the respiratory virus under investigation **(Fig 3)**. This is a disadvantage when different viruses are collected from the air simultaneously. With semi-solid gelatin, infectious pH1N1 virus and HMPV were equally well collected. In addition, with semi-solid gelatin, a correct jet-to-plate distance is maintained for accurate size fractionation of aerosols and droplets, and is also sufficiently solid for it to be safely transported, which is an enormous advantage over VTM.

When the collection efficiency of the three air samplers for different respiratory viruses was compared, the BioSampler performed best by collecting infectious virus and virus RNA of all four viruses with high efficiency. Also the cascade impactor collected high amounts of infectious virus, however, the amount of virus captured in each stage varied for the different respiratory viruses. The largest difference was observed in stage 6 of the cascade impactor, where substantial lower amounts of HMPV, PIV3 and RSV were collected as compared to pH1N1 virus, suggesting that fewer virus particles of the Paramyxo- and Pneumoviridae were contained in aerosols of size 1.1 – 0.6 um.. A possible explanation for this observation might be the pleomorphic character, size differences or aggregation of these viruses ^43,44^. Despite the fact that influenza viruses also form filamentous virus particles, it has been shown that passaging the viruses on eggs and cells results in the formation of mainly spherical particles of around 200 nm ^45,46^. Although the influenza strain used here was a low-passage clinical isolate, the ratio of spherical and filamentous particles in the influenza virus stock is not known, also not for the HMPV, PIV3 and RSV stocks.

The BioSampler and cascade impactor have a cut-off size of approximately 300 and 650 nm, respectively, meaning that only particles larger than the cut-off size are collected with high efficiency. Therefore, to collect smaller aerosols more efficiently, an electrostatic precipitator was developed in-house and also tested with all four respiratory viruses. Unfortunately, its overall performance was disappointing as only very low amounts of infectious virus and viral RNA was collected. A possible reason may be that droplets and aerosols were insufficiently charged with cations resulting in droplets and aerosols moving with the air through the sampler, rather than precipitating in the collection medium.

Several other studies have also evaluated the collection efficiency of the BioSampler and cascade impactor, or have also employed electrostatic precipitation to collect microorganisms from air ^42,47–49^. However, experimental set-ups including nebulizer type, collection medium and the applied flow rate are not uniform among studies. Particularly in the case of electrostatic precipitation no device is commercially available yet and hence air samplers are custom made designs ^20,23,50,51^. This makes it very difficult to directly compare the performance of the air samplers evaluated here, with other studies.

In conclusion, in the present study, the commercially available BioSampler and cascade impactor are both capable of collecting respiratory viruses while maintaining their infectivity during the sampling process. With the cascade impactor quantitative data on the sizes of virus-containing particles can be obtained, and in combination with a semi-solid gelatin layer as collection surface, the cascade impactor is also easy to use in various field settings such as hospitals. With the BioSampler size fractionation of the collected aerosols and droplets is unfortunately not possible. However, collection is more facile, since only one air sample is obtained per collection moment and no post air sampling processing is needed. The choice for either of the two air samplers therefore also depends on the environment in which it is to be used and on the research questions to be addressed. Overall, implementation of these air samplers in field studies will help to obtain more quantitative data on the amount of infectious respiratory virus that is present in the air, thereby generating a better understanding of respiratory virus transmission.

## Data availability

All data are available from the corresponding author (S.H.) on reasonable request.

## Acknowledgements

This work was financed through an NWO VIDI grant (contract number 91715372), NIH/NIAID contract HHSN272201400008C and European Union’s Horizon 2020 research and innovation program VetBioNet (grant agreement No 731014).

